# Generalized mutualisms promote range expansion in both plant and ant partners

**DOI:** 10.1101/2023.04.12.536632

**Authors:** Pooja Nathan, Evan P. Economo, Benoit Guénard, Anna Simonsen, Megan E. Frederickson

## Abstract

Mutualism improves organismal fitness, but strong dependence on another species can also limit a species’ ability to thrive in a new range if its partner is absent. We assembled a large, global dataset on mutualistic traits and species ranges to investigate how multiple plant-animal and plant-microbe mutualisms affect the spread of legumes and ants to novel ranges. We found that generalized mutualisms increase the likelihood that a species establishes and thrives beyond its native range, whereas specialized mutualisms either do not affect or reduce non-native spread. This pattern held in both legumes and ants, indicating that specificity between mutualistic partners is a key determinant of ecological success in a new habitat. Our global analysis shows that mutualism plays an important, if often overlooked, role in plant and insect invasions.

## Introduction

### Life did not take over the globe by combat, but by networking

Margulis & Sagan (1986) When a species arrives in a new range, its ecological success in the novel habitat is determined by biotic and abiotic factors (Paquette and Hargreaves 2021). Past studies have investigated how climate, life history, and antagonistic species interactions (e.g., enemy release, Meijer *et al*. 2016) influence range expansions and biological invasions. Fewer studies investigate how positive interactions such as mutualisms shape range dynamics (Shaw *et al*. 2021; Moles *et al*. 2022; Fowler *et al*. 2023). Here, we ask: how does mutualism contribute to the global spread of legumes and ants?

How mutualism affects the eco-evolutionary success of participating species is the subject of debate. Mutualism increases fitness or population size by helping organisms overcome resource limitation, cope with stressors, or avoid or escape enemies. Thus, mutualism might increase the niche breadth or geographic range of species (Bruno *et al*. 2003; Afkhami *et al*. 2014a; Batstone *et al*. 2018), buffer against environmental change (Lau & Lennon 2012), or even create the kind of ecological opportunity that leads to adaptive radiation (Weber & Agrawal 2014; Zeng & Wiens 2021). However, mutualism may also make populations more vulnerable to environmental change and result in local or global co-extinctions (Brodie *et al*. 2014). If one species is highly specialized on another, it may struggle to persist under new environmental conditions or in a new range if its mutualistic partner becomes scarce, even if there are no direct negative effects of the new environment on the focal species (Brodie *et al*. 2014, Kiers *et al*. 2015, Benning & Moeller 2021). Moreover, mutualisms are expected to break down because of the threat of ‘cheating’ or exploitation (Porter & Simms 2014; Jones *et al*. 2015, but see Frederickson 2017) and widespread context-dependency (Chamberlain *et al*. 2014). Such ecological or evolutionary fragility might prevent mutualists from thriving in new habitats or environments (Kiers *et al*. 2010), especially if mutualisms are most beneficial under the historical abiotic or biotic conditions in which the interaction evolved (Batstone *et al*. 2020).

Recent work has found evidence for both views: some studies have found that mutualism facilitates niche and range expansion (Afkhami *et al*. 2014a), while other studies have found the opposite (Duffy & Johnson 2017; Simonsen *et al*. 2017; Benning & Moeller 2021). One key determinant of the direction of effects appears to be the degree of specialization of the mutualism (Batstone *et al*. 2018; Harrison *et al*. 2018; Parshuram *et al*. 2023). Typically, the more generalized an interaction, the more likely a species is to encounter a compatible mutualist in its new range. If finding compatible partners in the new range is likely, and benefits derived from mutualism ameliorate abiotic or biotic stressors, mutualism can promote range expansion by allowing hosts to survive in otherwise uninhabitable ranges (Afkhami *et al*. 2014a). In contrast, if compatible partners are scarce and the species requires specific mutualistic partners, being mutualistic may hinder range expansion (Duffy & Johnson 2017; Simonsen *et al*. 2017; Harrison *et al*. 2018). Thus, specialization may constrain a mutualistic species’ ecological success outside its native range (Harrison *et al*. 2018).

These ideas can be extended to the study of how species introduced to new habitats by anthropogenic activity come to colonize them successfully—i.e., to invasion biology. Indeed, there is growing evidence that successful plant invaders have generalized interactions with pollinators, seed dispersers, and arbuscular mycorrhizal fungi (Richardson *et al*. 2000; Albrecht *et al*. 2014; García *et al*. 2014; Menzel *et al*. 2017). Furthermore, mutualism is predicted to cause “invasional meltdown” (Simberloff & von Holle 1999), whereby positive feedback between mutualistic co-invaders amplifies their spread or impacts (O’Dowd *et al*. 2003; Prior *et al*. 2014).

We investigated mutualism’s effects on introduced range size in ants and legumes. We chose these taxa because many ants and legumes are highly invasive species; these two families alone constitute 11% of the taxa on the IUCN’s list of the World’s 100 Worst Invasive Alien Species (“GISD” 2022). Substantial prior research has explored ecological factors that influence ant and legume invasions separately (e.g., Bradshaw *et al*. 2008; Wong *et al*. 2023), but no study has systematically evaluated the contribution of many mutualism types to the global spread of ants or legumes.

We investigated how ant-plant and plant-microbe mutualisms affect ant and legume range expansions. Mutualistic ants provide protection against enemies or seed dispersal services to many legumes, which reward ants with extrafloral nectar (EFN), nest sites in the form of live plant cavities (domatia), or lipid-rich appendages on seeds (elaiosomes). Most, but not all, legumes also engage in nutritional mutualisms with nitrogen-fixing rhizobia and phosphorus-provisioning mycorrhizae. These mutualisms span a continuum from highly generalized interactions such as seed dispersal by ants (myrmecochory) and ant-plant protection mutualisms mediated by EFNs, to highly specialized symbioses between ants and domatia-bearing plants (myrmecophytes), with variable partner specificity within legume-rhizobium and legume-mycorrhiza mutualisms. We compiled global range and trait datasets to test how mutualisms affect the invasive spread of legumes and ants in what is, to our knowledge, the largest comprehensive assessment to-date of mutualism and range size.

## Materials and methods

### Legume range data

We obtained range size data from a previous global analysis of legume geographic distributions (Simonsen *et al*. 2017). This dataset includes range data for over 3500 legume species and was originally compiled from records in the International Legume Database & Information Service (ILDIS, www.ildis.org). ILDIS lists which countries, states, or provinces each legume species occurs in, and whether it is native or introduced in each politically defined region. Summing the areas of each region where a legume occurs over-estimates range size because a species may occupy only a small portion of a politically defined spatial polygon. Therefore, we followed previous studies (Simonsen *et al*. 2017; Harrison *et al*. 2018; Parshuram *et al*. 2023) and analyzed the number of non-contiguous introduced ranges for each species, rather than total introduced range area.

### Legume mutualistic traits

We obtained data on legume associations with rhizobia from Simonsen *et al*. (2017) who compiled it from Werner *et al*. (2014); this study scored over 5000 legume species as ‘symbiotic’ or not, based on whether the legume forms nodules with rhizobia. We obtained data on interactions with mycorrhizae from an extensively curated dataset on the mycorrhizal associations of 14,870 plant species (Soudzilovskaia *et al*. 2020). We obtained data on the presence of EFNs from a curated, and frequently updated global list of plant species with EFNs (www.extrafloralnectaries.org) (Weber *et al*. 2015). Fabaceae has more EFN-producing plant species than any other plant family (Weber *et al*. 2015). We used data on which legumes have ant domatia from Chomicki and Renner (2015). Because these published lists of EFN-and domatia-bearing plants contain most records of EFN-and domatia-bearing plants known to science, we assumed that domatia and EFNs are absent from legumes not listed in Chomicki and Renner (2015), and www.extrafloralnectaries.org respectively. However, it is possible that this approach mis-classifies some legume species that are not well-studied. To control for this effect, we included the number of publications for each legume species available through CrossRef as a covariate. In contrast, for the legume-rhizobium and legume-mycorrhiza mutualisms, if there was no data for a legume species in Simonsen *et al*. (2017) or Soudzilovskaia *et al*. (2020), respectively, we omitted the species from further analysis, since our data on these mutualisms was not exhaustive. We harmonized species names across datasets and used the R package *taxize* (Chamberlain & Szöcs 2013) to verify accurate species names and synonyms. The final dataset consisted of all legume species for which we had range size data and some mutualistic trait data.

### Ant range data

We obtained data on the native and introduced range sizes of ants from the Global Ant Biodiversity Informatics (GABI) Project (Guénard *et al*. 2017). To calculate the total area of the native and introduced range for each species, we summed the areas of the spatial polygons making up an ant species’ native or introduced range. This approach over-estimates ant range size, so again we calculated the number of non-contiguous introduced ranges for each ant species by determining whether all pairs of spatial polygons making up its introduced range share a border using the *st_disjoint* function in the *sp* package in R (Pebesma *et al*. 2023) and converting the resulting matrix into a graph using the *igraph* package (Nepusz *et al*. 2023). We counted the independent subgraphs in the graph to determine the number of non-contiguous introduced ranges.

### Ant mutualistic traits

We obtained data on which ant species visit EFNs, nest in domatia, and disperse elaiosome-bearing seeds from Kaur *et al*. (2019) where the authors text-mined the abstracts of nearly 90,000 published articles about ants to algorithmically score which ant species do or do not visit EFNs, nest in domatia, or disperse seeds. For seed dispersal, they validated the text-mining results by comparing to a global list of seed-dispersing ants assembled and curated by a myrmecochory expert (Dr. Crisanto Gomez). We treated these ant traits (visits EFNs, nests in domatia, disperses seeds) as binary, categorial variables.

### Data analysis

All analyses were conducted in R version 4.2.0 (R Core Team 2022). For legume and ant introductions, we fit two models each: a generalized linear mixed model with a binomial error distribution for whether (1) or not (0) a species has been successfully introduced outside its native range, and a linear mixed model with a Gaussian error distribution of the log-transformed number of introduced ranges for species with at least one non-native range in the dataset (i.e., the second model excluded species that have not established outside their native range). For legume range size, we had two datasets that we modelled separately: one had trait data on the presence or absence of EFNs, domatia, and nodules for 3977 legume species, and the other had trait data on the presence or absence of AM and EM fungi for 723 legume species. For the larger dataset, we included the main effects of EFNs, domatia, and nodules and all possible interactions as predictors but simplified models by removing non-significant interactions. For the mycorrhizal dataset, we did not have sufficient degrees of freedom to model all five mutualistic traits and their interactions, so we fit a model with AM fungi, EM fungi, and their interaction as predictors, and then omitted the non-significant interaction and re-fit. Using the smaller mycorrhizal dataset, we also fit models with a count of the number of mutualisms each legume species engages in as a continuous variable (range: 0-4, as no legume has EFNs, domatia, nodules, AM, and EM; see SI). In all legume models, we included several other predictors available from Simonsen *et al*. (2017)’s dataset, specifically plant life history (annual or perennial), plant habit (woody or herbaceous), the number (range 0-9) of distinct human uses and the absolute latitude of the centroid of each legume’s native range. We also included counts of number of publications available for each legume species, determined using the *cr_works* function in the *rcrossref* package (Chamberlain *et al*. 2020) as a predictor. For ant range size, we fit models with EFN visitation, residing in domatia, and seed dispersal as predictors. We calculated the absolute latitude of the centroid of each ant’s native range and included it as a covariate in all ant models. There is no equivalent in the ant range dataset to the ‘number of human uses’ variable in the legume range dataset, but we used available information on the mode of ant introduction to classify introduced ants as “intercepted” (i.e., at customs), “indoor introduced,” or “exotic (naturalized)” as per Wong *et al*. (2023), and tested whether mutualistic traits affected the probability that an ant would be introduced indoors, intercepted by customs, or become naturalized in its introduced range. We used the *lmer* and *glmer* functions to fit models (package *lme4*; Bates 2010) and then ran Type III Wald chi-square tests to assess statistical significance using the *Anova* function from the *car* package (Fox *et al*. 2015). In both legume and ant models, we accounted for the phylogenetic non-independence of species in our dataset (Felsenstein 1985) by including tribe as a random effect in the mixed models. We also fit phylogenetic generalized least squares (PGLS) models (methods in SI), although they have lower statistical power because they exclude species for which we have trait and range data but no phylogenetic information.

## Results

### Legumes

In total, the legume range dataset included 3977 species, of which 3581 associate with nitrogen-fixing bacteria, 280 have extrafloral nectaries, and 24 have domatia. The smaller dataset on legumes with mycorrhizal fungi included 572 with arbuscular mycorrhizal (AM) fungi, 38 with ectomycorrhizal (EM) fungi, 80 with both AM and EM fungi (690 total taxa with mycorrhizae), and 33 without mycorrhizal fungi.

The presence of EFNs, but not domatia, nodules, or either type of mycorrhizae, significantly increased the likelihood of a legume successfully establishing in at least one non-native range (Figure 1, Table 1). Legumes with EFNs have also colonized significantly more introduced ranges than legumes without EFNs (Figure 1, Table 1). Although having domatia did not affect whether a legume species is introduced to at least one new range, domatia did significantly reduce the number of introduced ranges (although there were only 7 introduced legumes with domatia; Figure 1, Table 1). None of the belowground mutualisms significantly predicted introduction success or the number of introduced ranges (Figure 1, Table 1), although we observed the same trend reported by Simonsen *et al*. (2017), in which legumes that associate with rhizobia colonized fewer introduced ranges than non-nodulating legumes. The PGLS model also found a significant positive effect of EFNs, but no other mutualisms, on legume introductions (details in SI). Thus, overall, EFNs had consistently large, positive effects on legume introductions, domatia had weaker negative to neutral effects, and mycorrhizal fungi and nodules had no significant effects. Nonetheless, there was a significant, positive effect of engaging in more mutualisms on legume introduction success, and a marginally significant (p = 0.07) positive effect of engaging in more mutualisms on the number of introduced ranges (Figure S1, Table S1). Nodulation and AM fungi were the most widespread mutualisms in the legume dataset (additional details in SI) and this result suggests they may increase introduction success when combined with other mutualistic traits, but we did not fit a model testing all five mutualistic traits (EFNs, domatia, nodules, and AM and EM fungi) and all their possible interactions because of the small size of the legume-mycorrhizae dataset.

**Figure 1.**
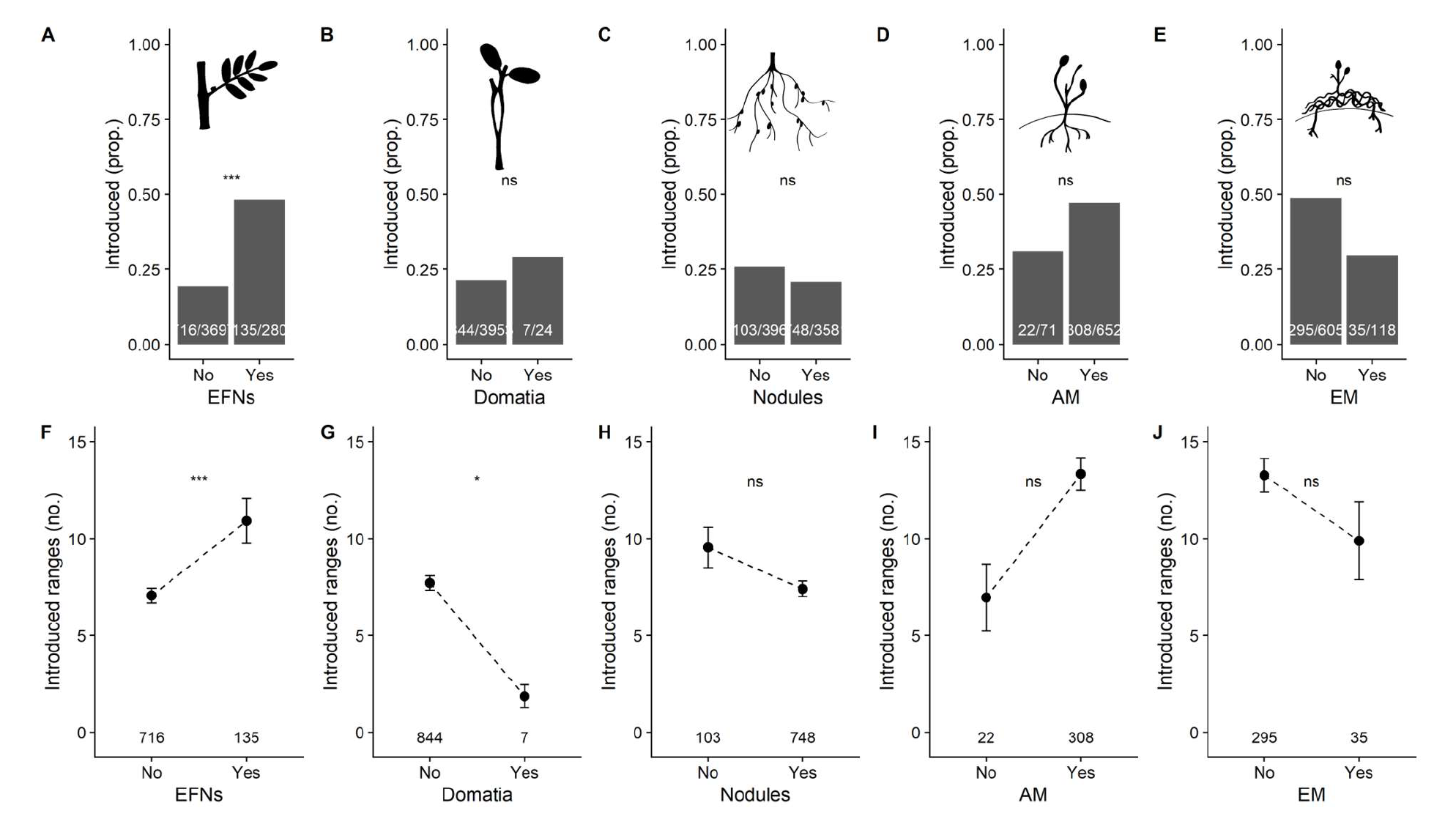
Relationship between the presence or absence of EFNs (A, F), domatia (B, G), nodules (C, H), arbuscular mycorrhizae (D, I), or ectomycorrhizae (E, J) and the proportion of introduced species (top) or number of introduced ranges (bottom, mean ± 1SE). Above the x-axis is the proportion of introduced taxa/total taxa (top) or number of introduced species (bottom) in each category. Significant effects indicated by *** for *p* ≤ 0.001, ** for *p* ≤ 0.01, and * for *p* ≤ 0.05. See Table 1 for complete results of statistical models.

**Table 1.** Model results for the effects of A. EFNs, domatia, and nodules and B. AM and EM fungi on whether legumes are successfully introduced to any novel range (left columns), and if successfully introduced, how many introduced ranges they spread to (right columns). Interactions among mutualistic traits were never significant and omitted from final models. Significant effects *italicized* and indicated by *** for *p* ≤ 0.001, ** for *p* ≤ 0.01, and * for *p* ≤ 0.05

| **A.** | **Successful introduction? (Binomial GLMM)** |  | **Number of introduced ranges (Gaussian LMM)** |  |
| --- | --- | --- | --- | --- |
| **Fixed effects** | **Estimate** | **SE** | **Estimate** | **SE** |
| Intercept | *-3.544\*\*\** | *0.296* | *0.850\*\*\** | *0.163* |
| EFNs? | *1.254\*\*\** | *0.195* | *0.342\*\*\** | *0.099* |
| Domatia? | 0.431 | 0.642 | *-0.986\** | *0.384* |
| Nodules? | 0.399 | 0.238 | -0.208 | 0.136 |
| Total native area | *0.186\*\** | *0.057* | -0.051 | 0.037 |
| Annual? | *0.670\*\*\** | *0.186* | 0.010 | 0.120 |
| Woody? | *0.784\*\*\** | *0.162* | *-0.218\** | *0.100* |
| Latitude | 0.091 | 0.069 | -0.064 | 0.042 |
| Number of human uses | *1.002\*\*\** | *0.049* | *0.320\*\*\** | *0.016* |
| Publication count | 0.064 | 0.048 | 0.022 | 0.032 |
| **Random effect** | **Variance** |  | **Variance** |  |
| Tribe | 0.511 |  | 0.040 |  |

| **B.** | **Successful introduction? (Binomial GLMM)** |  | **Number of introduced ranges (Gaussian LMM)** |  |
| --- | --- | --- | --- | --- |
| **Fixed effects** | **Estimate** | **SE** | **Estimate** | **SE** |
| Intercept | *-2.465\*\*\** | *0.535* | *0.816\*\*\** | *0.266* |
| AM? | -0.164 | 0.391 | 0.227 | 0.224 |
| EM? | -0.226 | 0.377 | -0.023 | 0.218 |
| Total native area | 0.240 | 0.145 | *-0.178\*\** | *0.064* |
| Annual? | 0.699 | 0.512 | 0.193 | 0.235 |
| *Woody?* | *0.911\** | *0.360* | *-0.367\** | *0.165* |
| Latitude | 0.201 | 0.151 | -0.116 | 0.070 |
| Number of human uses | *0.951\*\*\** | *0.087* | *0.315\*\*\** | *0.025* |
| Publication count | 0.197 | 0.130 | 0.030 | 0.055 |
| **Random effect** | **Variance** |  | **Variance** |  |
| Tribe | 0.779 |  | 0.096 |  |

As in Simonsen *et al*. (2017), legumes with more human uses were both more likely to be introduced and established in more introduced ranges (Table 1), and annuals were significantly more likely to be introduced than perennials, although only in the model of EFN, domatia, and nodulation effects (Table 1). Native range area had inconsistent effects across models; legumes with larger native range areas were more likely to be introduced in the model of EFN, domatia, and nodulation effects (Table 1), but in the smaller mycorrhizal dataset, models predicted that introduced legumes with larger native range areas have spread to fewer introduced ranges (Table 1). Woody legumes were significantly more likely to be introduced but spread to fewer introduced ranges than herbaceous legumes in both the model of EFN, domatia, and nodulation effects, and in the smaller mycorrhizal dataset (Table 1). The absolute midpoint latitude of a legume’s native range never significantly predicted its introduction success or number of introduced ranges, meaning there was no difference among tropical, sub-tropical, and temperate legumes in non-native spread. The number of studies on each species, a proxy for how well-studied a species is, also never significantly predicted introduction success or introduced range size in legumes.

Mutualistic traits generally had non-significant effects on the total native range area of legumes, except having both domatia and EFNs strongly decreased native range area (Figure S2, Table S2). However, only four legume species in the dataset have domatia and EFNs (three ant-acacias and *Humboldtia laurifolia*). Covariate effects on native range size are described in the SI.

### Ants

In total, we had range and trait data for 3023 species, of which 115 visit EFNs, 297 disperse seeds, and 58 nest in domatia. Ant species that visit EFNs were more likely to be introduced and have spread to more introduced ranges than other ants (Figure 2, Table 2). Seed-dispersing ant species were also more likely to be introduced but have not spread to more introduced ranges than ants that do not disperse seeds (Figure 2, Table 2). In contrast, nesting in domatia had no significant effect on introduction success or number of introduced ranges (Figure 2, Table 2). Ants with larger native range areas were more likely to be introduced but have not invaded more ranges (Table 2). The midpoint latitude of an ant’s native range was significantly negatively associated with both introduction success and the number of introduced ranges (Table 2), meaning that tropical ants have had greater non-native spread than temperate ant species. The PGLS model also found significant, positive effects of visiting EFNs and dispersing seeds, but not nesting in domatia, on the number of introduced ranges (details in SI).

**Figure 2.**
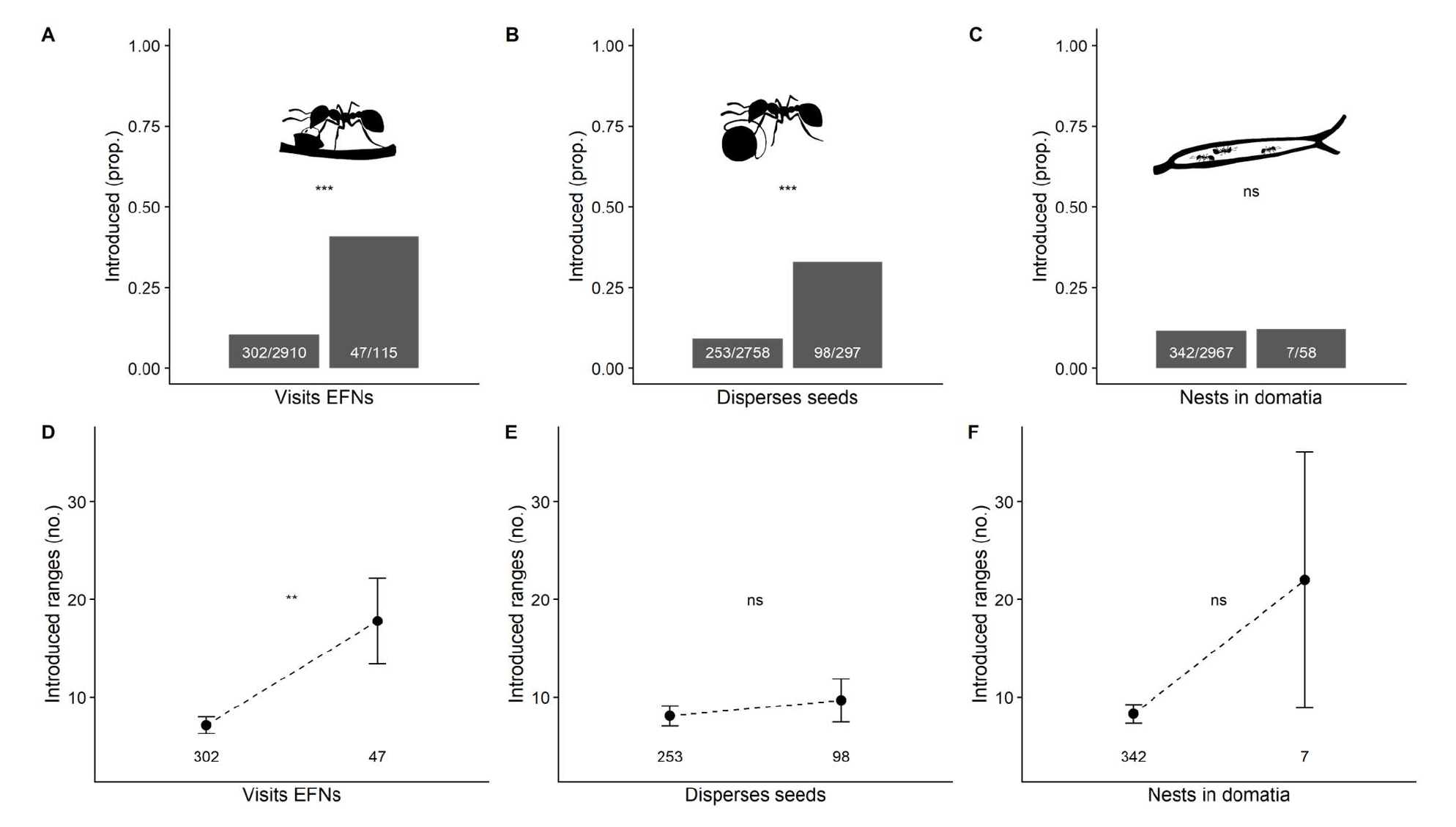
Relationship between ants visiting EFNs (A, D), dispersing seeds (B, E), and nesting in domatia (C, F) and the proportion of introduced species (top) or introduced range area (bottom, mean ± 1SE). Above the x-axis is the proportion of introduced taxa/total taxa (top) or number of introduced species (bottom) in each category. Significant effects indicated by *** for *p* ≤ 0.001, ** for *p* ≤ 0.01, and * for *p* ≤ 0.05. See Table 2 for complete results of statistical models.

**Table 2.**
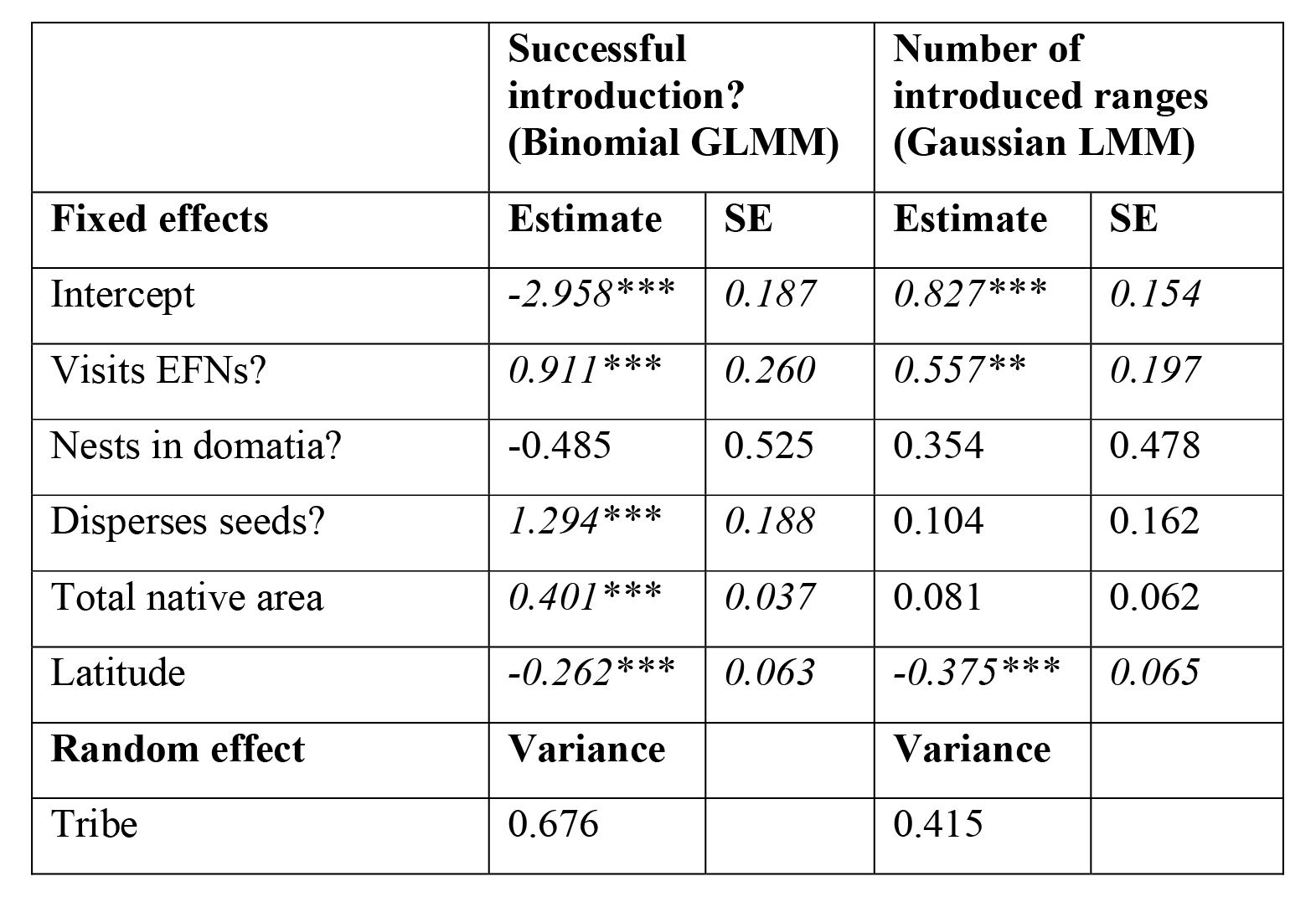
Model results for the effects of visiting EFNs, dispersing seeds, and nesting in domatia on whether ants are successfully introduced to any novel range (left columns), and if successfully introduced, their introduced range area (right columns). Interactions among mutualistic traits were never significant and omitted from final models. Significant effects *italicized* and indicated by *** for *p* ≤ 0.001, ** for *p* ≤ 0.01, and * for *p* ≤ 0.05.

Mutualistic traits never had significant interaction effects on introduction success or introduced range area in the mixed models, so the final models fit main effects of mutualistic traits only (Table 2). In other words, mutualisms had additive effects on ant introductions, and not sub-additive or synergistic effects. There was also a significantly positive effect of the number of mutualisms on ant introduction success and number of introduced ranges (Figure S3, Table S3).

Unlike in legumes, mutualisms significantly predicted ant native range sizes. Ants that visit EFNs or disperse seeds had significantly larger native range areas than species that do not participate in these mutualisms (Figure S4, Table S4, and see SI for covariates).

Ants that visit EFNs were significantly more likely to establish indoors, but no more likely to be naturalized outdoors or intercepted at customs than other ants (Figure 3, Table S5). Nesting in domatia had no significant effects on customs interception, indoor establishment, or naturalization (Figure 3, Table S5). Ants that disperse seeds were significantly more likely to be intercepted at customs, but significantly less likely to establish indoors, and equally likely to be naturalized as other ants (Figure 3, Table S5).

**Figure 3.**
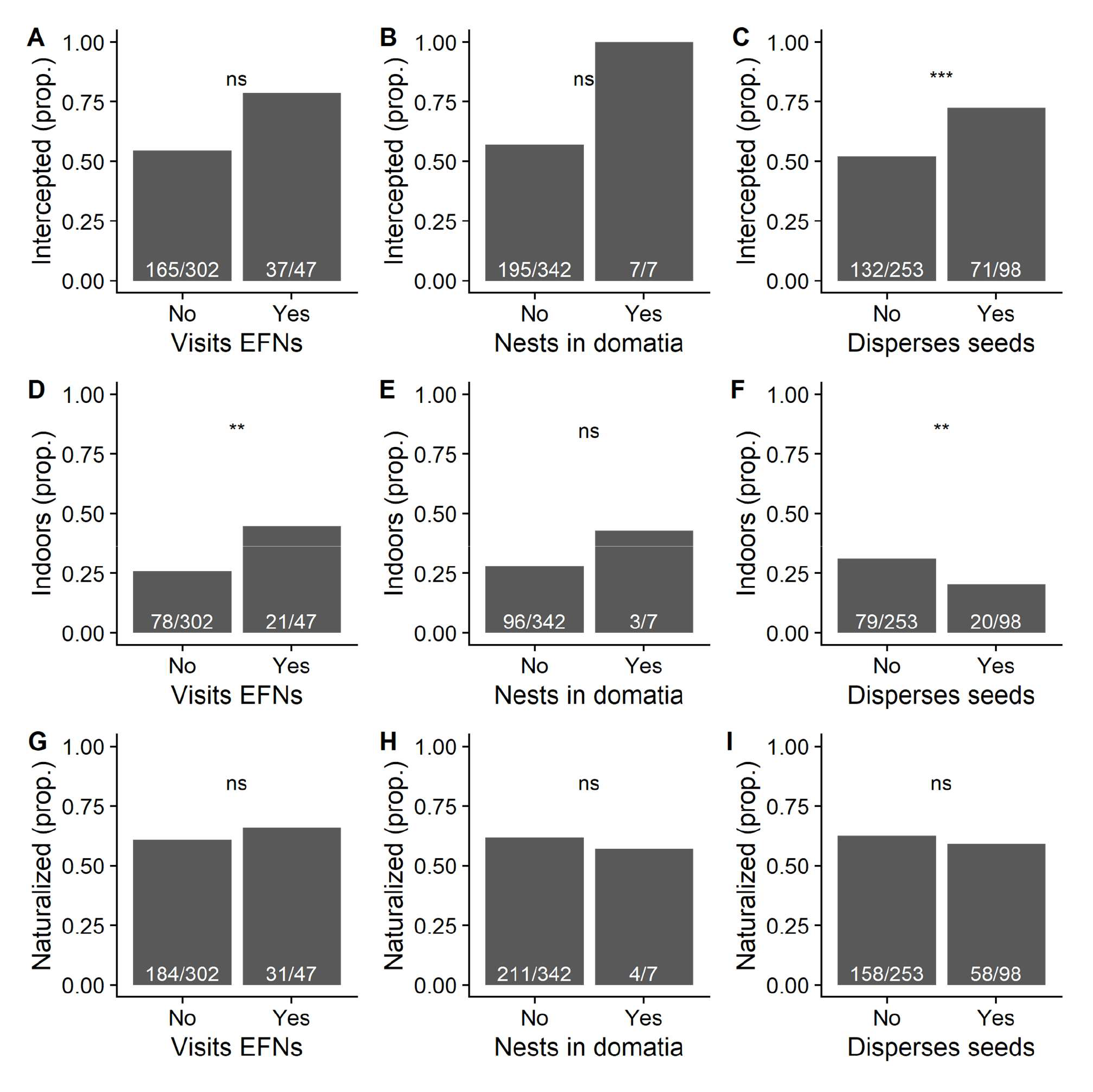
Relationship between ants visiting EFNs, dispersing seeds, and nesting in domatia and whether an ant species is intercepted at customs (top), establishes indoors (middle), or have become naturalized outdoors (bottom) in non-native ranges. Numbers above the x-axis indicate the number of intercepted/indoor/naturalized taxa/total introduced taxa. Significant effects indicated by *** for *p* ≤ 0.001, ** for *p* ≤ 0.01, and * for *p* ≤ 0.05. See Table S6 for complete results of statistical models.

## Discussion

Several mutualisms affected introduction success and invasive spread, with generalized mutualisms facilitating range expansion. Notably, the very generalized mutualism between EFN-bearing legumes and EFN-visiting ants increased the likelihood of being introduced to new ranges, as well as the number of introduced ranges, in both partners. Dispersing seeds, another generalized mutualism, was also associated with increased invasiveness in ants. In contrast, associating with rhizobia or mycorrhizal fungi did not significantly affect the non-native spread of legumes, and residing in domatia had no significant effect on ant invasions. Finally, producing domatia constrained non-native spread in legumes. These results suggest that generalized mutualisms improve the likelihood of successful colonization of new ranges, with specialized mutualisms having no, or opposite, effects.

There is debate about how mutualisms impact ecological success. On the one hand, associating with a mutualist may enable a species to better tolerate abiotic or biotic stressors in a new range (Afkhami *et al*. 2014b; Batstone *et al*. 2018). On the other hand, specialization on a mutualistic partner may reduce a species’ ability to colonize ranges where its partner is absent (Simonsen *et al*. 2017; Harrison *et al*. 2018b; Benning & Moeller 2019; but see Birnbaum *et al*. 2012). Our study suggests that both scenarios occur, although there was stronger evidence for the former. Very generalized mutualisms may allow species to associate with previously unseen mutualistic partners and colonize new ranges (Richardson *et al*. 2000).

Our study is the first to find that the mutualism between EFN-bearing plants and EFN-visiting ants has reciprocally positive effects on range expansions in both partners observable at the macroecological scale. In all models, legumes with EFNs were better invaders, as were ants that visited EFNs. The mutualism between EFN-visiting ants and EFN-bearing plants is usually generalized in both plant and ant partners (Heil & McKey 2003; Marazzi *et al*. 2013; but see Heil *et al*. 2010). EFN-visiting ants tend to be cosmopolitan, and EFN-producing legume species may have higher ecological success because they recruit local ant species, some of which may serve as bodyguards against herbivores in new ranges, even without sharing evolutionary history. This is evidenced by records of native ant species visiting the EFNs of invasive plant species (Eichhorn *et al*. 2011; Johnson *et al*. 2019). Our results also suggest that ant species that visit EFNs are more successful at colonizing new habitats. Extrafloral nectar from native plant species may serve as a valuable food resource for newly introduced ant species (as recorded by Ness & Bronstein 2004; Lach *et al*. 2009; Savage *et al*. 2009; Riginos *et al*. 2015) even when the species have not interacted before. There are also many reports of invasive ants visiting EFNs on invasive plants (Lach *et al*. 2010; Eichhorn *et al*. 2011; Gippet *et al*. 2018), even when they come from different regions, suggesting that little or no coevolution is required for a new interaction to be established. In contrast, the mutualism between domatia-nesting ants and domatia-bearing plants is quite specialized in both ant and plant partners (Chomicki and Renner 2015; but see Chanam *et al*. 2014; Volp *et al*. 2022), likely explaining why domatia reduced invasive spread in legumes, while nesting in domatia had no effect in ant introductions.

Much as ants visit EFNs on unfamiliar plants, seed-dispersing ants often pick up elaiosome-bearing seeds from invasive plant species even if they have not encountered them before (Passos & Oliveira 2003; Rowles & O’Dowd 2009; Stuble *et al*. 2010; Rodriguez-Cabal *et al*. 2012; Devenish *et al*. 2019; Anjos *et al*. 2020) and as expected, seed-dispersing ants had larger native and introduced range sizes than ants that do not pick up seeds. While seed-dispersing ants often interact with many plant species, so long as their seeds have elaiosomes, this interaction may be less generalized from the plant perspective, with only a few ant species being effective seed dispersers (Manzaneda & Rey 2009; Ness *et al*. 2009). Thus, while ants may benefit from “dispersing” seeds in their new range, the same cannot be assumed for introduced myrmecochorous plants (Warren *et al*. 2010). However, we were limited by the availability of trait data to testing the effects of the mutualism between seed-dispersing ants and myrmecochorous plants only in ants, not in legumes.

In legumes, associating with rhizobia had no significant negative effects on introduction success. A previous study using the same dataset (Simonsen *et al*. 2017) found that legumes that form nodules with rhizobia have spread to fewer new ranges than non-nodulating legumes, and we found a non-significant trend in the same direction, although our model structures were different. Our analysis included more mutualistic traits than Simonsen *et al*. (2017), and we had limited statistical power to test for all the possible interactions between mutualistic traits and covariates. In Simonsen *et al*. (2017), however, the higher non-native spread of non-symbiotic legumes was only observed only among species that had a low number of human uses. The ability to nodulate and hence fix nitrogen is a useful trait for humans (e.g., in forage or rotation crops), thus humans may deliberately introduce symbiotic legumes and overcome the establishment barrier caused by lack of rhizobial partners. In a follow-up study to Simonsen *et al*. (2017), Harrison *et al*. (2018) found that even within nodulating legumes, species that associate with more rhizobial strains have larger introduced ranges than those that interact with fewer strains, substantiating the trend of generalized mutualisms increasing range expansion that we observe across mutualism types, but within a single kind of association. Furthermore, Parshuram *et al*. (2023) recently found that specialization in legume-rhizobium mutualisms constrains range expansions only in polyploid, and not in diploid legumes. Thus, variation among nodulating legumes in their specificity on rhizobia and interactions between mutualism and other traits probably explain why nodulation alone did not predict legume introduction success or invasive spread in our analysis.

Interacting with AM fungi had a non-significant positive effect on the successful introduction of legumes, while interacting with EM fungi had a non-significant negative effect. The overwhelming majority of plant species are mycorrhizal (Brundrett 2009; Maherali *et al*. 2016) and thus our dataset contained only 33 legume species that did not partner with mycorrhizal fungi, meaning that we had low statistical power to detect effects of mycorrhizae on legume range expansions. Although we did not observe any significant effects of AM or EM fungi on legume introductions, the limiting effect of EM fungi on range expansions has been studied extensively in pines (Nuñez *et al*. 2009; Hayward *et al*. 2015; Policelli *et al*. 2019), which famously could not be successfully cultivated in the southern hemisphere until they were inoculated with soil from their native range containing suitable EM fungi (Richardson 2000).

In both legumes and ants, we found that participating in more mutualisms increased the likelihood of successful introduction and spread in the new range. This result suggests that the positive effects of generalized mutualisms counteract any negative effects of specialized mutualisms when a species colonizes a new range and that, overall, mutualism promotes, rather than hinders, range expansion. Recent studies have also shown that AM fungi and rhizobia act synergistically in their interactions with legumes, promoting plant growth when together more than expected from the growth-promotion effects of rhizobia or AM fungi alone (Larimer *et al*. 2014; Afkhami & Stinchcombe 2016; Afkhami *et al*. 2021). In legumes, the most common mutualisms are with rhizobia and AM fungi, and the synergistic effects of these mutualisms may drive the increase in colonization success we observe for legumes that participate in more types of mutualisms. Multiple mutualist effects (Afkhami *et al*. 2014) may provide more insights into the mechanisms of how multiple mutualistic partners influence each other and their shared host, but most current studies focus on only one kind of mutualism in isolation, especially at large biogeographic or macroevolutionary scales (Chamberlain & Rudgers 2012; Godschalx *et al*. 2015; Dutton *et al*. 2016; Afkhami *et al*. 2021; Laurich *et al*. 2022).

In legumes, mutualism had little effect on native range size, suggesting that different eco-evolutionary processes are involved in determining the native and introduced ranges of legumes (Sexton *et al*. 2009). The only strong effect of mutualism on legume native range size was the negative interaction between EFNs and domatia, in which legumes with both traits had significantly smaller native ranges than other legumes. This type of highly specialized ant-plant mutualism, in which plants both feed and house their ant symbionts, is rare, and occurred in only four legumes in our dataset. In contrast, in ants, mutualism had strong effects on both native and introduced ranges. Because the trait data for ants were generated by text-mining scholarly literature (Kaur *et al*. 2019), one explanation for this pattern is that mutualisms are more likely to be studied in widespread ant species and remain severely under-reported in ant species with small ranges. However, the text-mined trait dataset of Kaur *et al*. (2019) classified an ant species as a non-mutualist only if the species had received at least some study in the scholarly literature (i.e., they did not categorize unstudied ants, which might have smaller ranges on average, as non-mutualists). Furthermore, native range size is likely overestimated in our dataset (for both legumes and ants), because it is calculated from the summed areas of all politically defined spatial polygons where a species has historically occurred, regardless of how much of the polygon the species inhabits. We have no reason to suspect that native range size is overestimated more in mutualists than non-mutualists, but the native range results should nonetheless be interpreted with caution. This issue does not affect our analyses of introduction success or the number of introduced ranges, although both native and introduced ranges may still be under-estimated because of limited data on species occurrences, especially in the Global South.

Our results also suggest that human activity is an important determinant of legume and ant invasion success when viewed through a mutualism lens too. For legumes, the number of human uses contributed significantly to the likelihood of being introduced and the number of introduced ranges. Agriculture can be considered a mutualism between plants and humans (Boucher 1985; Spengler 2020; Angourakis *et al*. 2022) and the dispersal of cultivated species to new ranges is a consequence of this interaction. Reciprocally positive interactions between humans and legumes have clearly accelerated the introduction of legumes to new ranges.

Humans have also contributed to the non-native spread of ant species around the globe (Gippet *et al*. 2019; Gippet & Bertelsmeier 2021; Lach 2021). EFN-visiting ant species were more likely to establish indoors, possibly because of the abundance of sugary, nectar-like foods present in human settlements, because they were co-introduced with ornamental plants in indoor locations like greenhouses, or because they are more often tropical in origin (Luo *et al*. 2022) and thus thrive in warmer indoor environments. In contrast, seed-dispersing ants often originate from and invade mid-latitude regions, and they were more likely to be intercepted and less likely to be introduced indoors. The Mediterranean or temperate climates where myrmecochory is most common (Lengyel *et al*. 2010) may also be regions with strong customs surveillance of novel insect pests.

Mutualism is not generally considered a driver of macroecological patterns, but our results suggest that the benefits derived from generalized mutualisms increase ant and legume introductions globally. Indeed, mutualisms between introduced species may mediate the rapid spread of both partners, potentially accelerating biodiversity loss and other ecosystem changes, causing “invasional meltdown” (Simberloff & von Holle 1999; Traveset & Richardson 2014). Our global analysis suggests that the positive feedback that occurs between partners in mutualisms often amplifies the non-native spread of ants and legumes, but only when interactions are generalized.

## Acknowledgements

We acknowledge funding from the NSERC Discovery Grant to MEF.

## References

1. Afkhami, M.E., Friesen, M.L. & Stinchcombe, J.R. (2021). Multiple Mutualism Effects generate synergistic selection and strengthen fitness alignment in the interaction between legumes, rhizobia and mycorrhizal fungi. Ecol Lett, 24, 1824–1834.

2. Afkhami, M.E., McIntyre, P.J. & Strauss, S.Y. (2014a). Mutualist-mediated effects on species’ range limits across large geographic scales. Ecol Lett, 17, 1265–1273.

3. Afkhami, M.E., Rudgers, J.A. & Stachowitz, J.J. (2014c). Multiple mutualist effects: conflict and synergy in multispecies mutualisms. Ecology, 95, 833–844.

4. Afkhami, M.E. & Stinchcombe, J.R. (2016). Multiple mutualist effects on genomewide expression in the tripartite association between Medicago truncatula, nitrogen-fixing bacteria and mycorrhizal fungi. Mol Ecol, 25, 4946–4962.

5. Albrecht, M., Padrón, B., Bartomeus, I. & Traveset, A. (2014). Consequences of plant invasions on compartmentalization and species’ roles in plant–pollinator networks. Proceedings of the Royal Society B: Biological Sciences, 281.

6. Angourakis Id, A., Alcaina-Mateos, J., I.D., Madella, M. & Zurro, D. (2022). Human-Plant Coevolution: A modelling framework for theory-building on the origins of agriculture. PLoS One, 17, e0260904.

7. Anjos, D. v., Leal, L.C., Jordano, P. & Del-Claro, K. (2020). Ants as diaspore removers of non-myrmecochorous plants: a meta-analysis. Oikos, 129, 775–786.

8. Batstone, R.T., Carscadden, K.A., Afkhami, M.E. & Frederickson, M.E. (2018). Using niche breadth theory to explain generalization in mutualisms. Ecology, 99, 1039–1050.

9. Batstone, R.T., O’Brien, A.M., Harrison, T.L. & Frederickson, M.E. (2020). Experimental evolution makes microbes more cooperative with their local host genotype. Science (1979), 370.

10. Benning, J.W. & Moeller, D.A. (2019). Maladaptation beyond a geographic range limit driven by antagonistic and mutualistic biotic interactions across an abiotic gradient. Evolution (N Y), 73, 2044–2059.

11. Benning, J.W. & Moeller, D.A. (2021). Microbes, mutualism, and range margins: testing the fitness consequences of soil microbial communities across and beyond a native plant’s range. New Phytologist, 229, 2886–2900.

12. Birnbaum, C., Barrett, L.G., Thrall, P.H. & Leishman, M.R. (2012). Mutualisms are not constraining cross-continental invasion success of Acacia species within Australia. Divers Distrib, 18, 962–976.

13. Boucher, D. (1985). The Biology of Mutualism: Ecology and Evolution.

14. Bradshaw, C.J.A., Giam, X., Tan, H.T.W., Brook, B.W. & Sodhi, N.S. (2008). Threat or invasive status in legumes is related to opposite extremes of the same ecological and life-history attributes. Journal of Ecology, 96, 869–883.

15. Brodie, J.F., Aslan, C.E., Rogers, H.S., Redford, K.H., Maron, J.L., Bronstein, J.L., et al. (2014). Secondary extinctions of biodiversity. Trends Ecol Evol, 29, 664–672.

16. Bronstein, J. (Ed.). (2015). Mutualism.

17. Brundrett, M.C. (2009). Mycorrhizal associations and other means of nutrition of vascular plants: Understanding the global diversity of host plants by resolving conflicting information and developing reliable means of diagnosis. Plant Soil, 320, 37– 77.

18. Bruno, J.F., Stachowicz, J.J. & Bertness, M.D. (2003). Inclusion of facilitation into ecological theory. Trends Ecol Evol, 18, 119–125.

19. Chamberlain, S., Zhu, H., Jahn, N., Boettiger, C. & Ram, K. (2020). Client for Various “CrossRef” “APIs” [R package rcrossref version 1.2.0].

20. Chamberlain, S.A., Bronstein, J.L. & Rudgers, J.A. (2014). How context dependent are species interactions? Ecol Lett, 17, 881–890.

21. Chamberlain, S.A. & Rudgers, J.A. (2012). How do plants balance multiple mutualists? Correlations among traits for attracting protective bodyguards and pollinators in cotton (Gossypium). Evol Ecol, 26, 65–77.

22. Chamberlain, S.A. & Szöcs, E. (2013). taxize: taxonomic search and retrieval in R. F1000Res, 2, 191.

23. Chanam, J., Sheshshayee, M.S., Kasinathan, S., Jagdeesh, A., Joshi, K.A. & Borges, R.M. (2014). Nutritional benefits from domatia inhabitants in an ant–plant interaction: interlopers do pay the rent. Funct Ecol, 28, 1107–1116.

24. Chomicki, G. & Renner, S.S. (2015). Phylogenetics and molecular clocks reveal the repeated evolution of ant-plants after the late Miocene in Africa and the early Miocene in Australasia and the Neotropics. New Phytologist, 207, 411–424.

25. CRAN - Package igraph. (2023). . Available at: https://cran.r-project.org/web/packages/igraph/index.html. Last accessed 27 March 2023.

26. D, B. (2010). lme4: linear mixed-effects models using S4 classes. http://cran.r-project.org/web/packages/lme4/index.html.

27. Devenish, A.J.M., Gomez, C., Bridle, J.R., Newton, R.J. & Sumner, S. (2019). Invasive ants take and squander native seeds: implications for native plant communities. Biol Invasions, 21, 451–466.

28. Duffy, K.J. & Johnson, S.D. (2017). Specialized mutualisms may constrain the geographical distribution of flowering plants. Proceedings of the Royal Society B: Biological Sciences, 284.

29. Dutton, E.M., Luo, E.Y., Cembrowski, A.R., Shore, J.S. & Frederickson, M.E. (2016). Three’s a crowd: Trade-offs between attracting pollinators and ant bodyguards with nectar rewards in Turnera. American Naturalist, 188, 38–51.

30. Eichhorn, M.P., Ratliffe, L.C. & Pollard, K.M. (2011). Attraction of ants by an invasive Acacia. Insect Conserv Divers, 4, 235–238.

31. Felsenstein, J. (1985). Phylogenies and the comparative method. American Naturalist, 125, 1–15.

32. Fowler, J.C., Donald, M.L., Bronstein, J.L. & Miller, T.E.X. (2023). The geographic footprint of mutualism: How mutualists influence species’ range limits. Ecol Monogr, 93, e1558.

33. Fox, J., Weisberg, S., Adler, D., … D.B.-… : R.F. for & 2012, undefined. (2015). Package “car.” cran.uni-muenster.de.

34. Frederickson, M.E. (2017). Mutualisms Are Not on the Verge of Breakdown. Trends Ecol Evol, 32, 727–734.

35. García, D., Martínez, D., Stouffer, D.B. & Tylianakis, J.M. (2014). Exotic birds increase generalization and compensate for native bird decline in plant–frugivore assemblages. Journal of Animal Ecology, 83, 1441–1450.

36. Gippet, J.M., Liebhold, A.M., Fenn-Moltu, G. & Bertelsmeier, C. (2019). Human-mediated dispersal in insects. Curr Opin Insect Sci, 35, 96–102.

37. Gippet, J.M.W. & Bertelsmeier, C. (2021). Invasiveness is linked to greater commercial success in the global pet trade. Proc Natl Acad Sci U S A, 118, e2016337118.

38. Gippet, J.M.W., Piola, F., Rouifed, S., Viricel, M.R., Puijalon, S., Douady, C.J., et al. (2018). Multiple invasions in urbanized landscapes: interactions between the invasive garden ant Lasius neglectus and Japanese knotweeds (Fallopia spp.). Arthropod Plant Interact, 12, 351–360.

39. *GISD*. (2022). . Available at: http://www.iucngisd.org/gisd/100_worst.php. Last accessed 28 February 2022.

40. Godschalx, A.L., Schädler, M., Trisel, J.A., Balkan, M.A. & Ballhorn, D.J. (2015). Ants are less attracted to the extrafloral nectar of plants with symbiotic, nitrogen-fixing rhizobia. Ecology, 96, 348–354.

41. Guénard, B., Weiser, M.D., Gómez, K., Narula, N. & Economo, E.P. (2017). The Global Ant Biodiversity Informatics (GABI) database: Synthesizing data on the geographic distribution of ant species (Hymenoptera: Formicidae). Myrmecol News.

42. Harrison, T.L., Simonsen, A.K., Stinchcombe, J.R. & Frederickson, M.E. (2018a). More partners, more ranges: generalist legumes spread more easily around the globe. Biol Lett, 14.

43. Hayward, J., Horton, T.R., Pauchard, A. & Nunez, M.A. (2015). A single ectomycorrhizal fungal species can enable a Pinus invasion. Ecology, 96, 1438–1444.

44. Heil, M. & McKey, D. (2003). Protective Ant-Plant Interactions as Model Systems in Ecological and Evolutionary Research. Annual Review of Ecology, Evolution, and Systematics.

45. Heil, M., Orona-Tamayo, D., Eilmus, S., Kautz, S. & González-Teuber, M. (2010). Chemical communication and coevolution in an ant-plant mutualism. Chemoecology, 20, 63–74.

46. Johnson, L.R., Breger, B. & Drummond, F. (2019). Novel plant–insect interactions in an urban environment: enemies, protectors, and pollinators of invasive knotweeds. Ecosphere, 10, e02885.

47. Jones, E.I., Afkhami, M.E., Akçay, E., Bronstein, J.L., Bshary, R., Frederickson, M.E., et al. (2015). Cheaters must prosper: reconciling theoretical and empirical perspectives on cheating in mutualism. Ecol Lett, 18, 1270–1284.

48. Kaur, K.M., Malé, P.J.G., Spence, E., Gomez, C. & Frederickson, M.E. (2019). Using text-mined trait data to test for cooperate-and-radiate co-evolution between ants and plants. PLoS Comput Biol, 15, e1007323.

49. Kiers, E.T., Palmer, T.M., Ives, A.R., Bruno, J.F. & Bronstein, J.L. (2010). Mutualisms in a changing world: an evolutionary perspective. Ecol Lett, 13, 1459–1474.

50. Lach, L. (2021). Invasive ant establishment, spread, and management with changing climate. Curr Opin Insect Sci, 47, 119–124.

51. Lach, L., Hobbs, R.J. & Majer, J.D. (2009). Herbivory-induced extrafloral nectar increases native and invasive ant worker survival. Popul Ecol, 51, 237–243.

52. Lach, L., Tillberg, C. v. & Suarez, A. v. (2010). Contrasting effects of an invasive ant on a native and an invasive plant. Biol Invasions, 12, 3123–3133.

53. Larimer, A.L., Clay, K. & Bever, J.D. (2014). Synergism and context dependency of interactions between arbuscular mycorrhizal fungi and rhizobia with a prairie legume. Ecology, 95, 1045–1054.

54. Lau, J.A. & Lennon, J.T. (2012). Rapid responses of soil microorganisms improve plant fitness in novel environments. Proc Natl Acad Sci U S A, 109, 14058–14062.

55. Laurich, J.R., Reid, C.G., Biel, C., Wu, T., Knox, C. & Frederickson, M.E. (2022). The genetic architecture of multiple mutualisms and mating system in Turnera ulmifolia. bioRxiv, 2022.03.31.484762.

56. Lengyel, S., Gove, A.D., Latimer, A.M., Majer, J.D. & Dunn, R.R. (2010). Convergent evolution of seed dispersal by ants, and phylogeny and biogeography in flowering plants: A global survey. Perspect Plant Ecol Evol Syst, 12, 43–55.

57. Luo, Y., Taylor, A., Weigelt, P., Guénard, B., Economo, E.P., Nowak, A., et al. (n.d.). Climate and ant richness explain the global distribution of ant-plant mutualisms.

58. Maherali, H., Oberle, B., Stevens, P.F., Cornwell, W.K. & McGlinn, D.J. (2016). Mutualism Persistence and Abandonment during the Evolution of the Mycorrhizal Symbiosis. Am Nat, 188, E113–E125.

59. Manzaneda, A.J. & Rey, P.J. (2009). Assessing ecological specialization of an ant–seed dispersal mutualism through a wide geographic range. Ecology, 90, 3009–3022.

60. Marazzi, B., Bronstein, J.L. & Koptur, S. (2013). The diversity, ecology and evolution of extrafloral nectaries: current perspectives and future challenges. Ann Bot, 111, 1243– 1250.

61. Meijer, K., Schilthuizen, M., Beukeboom, L. & Smit, C. (2016). A review and meta-analysis of the enemy release hypothesis in plant-herbivorous insect systems. PeerJ, 2016, e2778.

62. Menzel, A., Hempel, S., Klotz, S., Moora, M., Pyšek, P., Rillig, M.C., et al. (2017). Mycorrhizal status helps explain invasion success of alien plant species. Ecology, 98, 92–102.

63. Moles, A.T., Dalrymple, R.L., Raghu, S., Bonser, S.P. & Ollerton, J. (2022). Advancing the missed mutualist hypothesis, the under-appreciated twin of the enemy release hypothesis. Biol Lett, 18, 20220220.

64. Nathan, P., Economo, E.P., Guénard, B., Simonsen, A. & Frederickson, M.E. (2023). Generalized mutualisms promote range expansion in both plant and ant partners. bioRxiv, 2023.04.12.536632.

65. Ness, J.H. & Bronstein, J.L. (2004). The Effects of Invasive Ants on Prospective Ant Mutualists. Biological Invasions 2004 6:4, 6, 445–461.

66. Ness, J.H., Morin, D.F. & Giladi, I. (2009). Uncommon specialization in a mutualism between a temperate herbaceous plant guild and an ant: are Aphaenogaster ants keystone mutualists? Oikos, 118, 1793–1804.

67. Nuñez, M.A., Horton, T.R. & Simberloff, D. (2009). Lack of belowground mutualisms hinders Pinaceae invasions. Ecology, 90, 2352–2359.

68. O’Dowd, D.J., Green, P.T. & Lake, P.S. (2003). Invasional ‘meltdown’ on an oceanic island. Ecol Lett, 6, 812–817.

69. Paquette, A. & Hargreaves, A.L. (2021). Biotic interactions are more often important at species’ warm versus cool range edges. Ecol Lett, 24, 2427–2438.

70. Parshuram, Z.A., Harrison, T.L., Simonsen, A.K., Stinchcombe, J.R. & contributed equally, A. (2022). Symbiosis with rhizobia limits range expansion only in polyploid legumes. bioRxiv, 2022.03.01.482489.

71. Passos, L. & Oliveira, P.S. (2003). Interactions between ants, fruits and seeds in a restinga forest in south-eastern Brazil. J Trop Ecol, 19, 261–270.

72. Pebesma, E., Bivand, R., Rowlingson, B., Gomez-Rubio, V., Hijmans, R. & Sumner, M. (2023). Classes and Methods for Spatial Data [R package sp version 1.6–0].

73. Policelli, N., Bruns, T.D., Vilgalys, R. & Nuñez, M.A. (2019). Suilloid fungi as global drivers of pine invasions. New Phytologist, 222, 714–725.

74. Porter, S.S. & Simms, E.L. (2014). Selection for cheating across disparate environments in the legume-rhizobium mutualism. Ecol Lett, 17, 1121–1129.

75. Prior, K.M., Saxena, K. & Frederickson, M.E. (2014). Seed handling behaviours of native and invasive seed-dispersing ants differentially influence seedling emergence in an introduced plant. Ecol Entomol, 39, 66–74.

76. Richardson, David.M. (Ed.). (2000). Ecology and Biogeography of Pinus.

77. Riginos, C., Karande, M.A., Rubenstein, D.I. & Palmer, T.M. (2015). Disruption of a protective ant–plant mutualism by an invasive ant increases elephant damage to savanna trees. Ecology, 96, 654–661.

78. Rodriguez-Cabal, M.A., Barrios-Garcia, M.N. & Nuñez, M.A. (2012). Positive interactions in ecology: filling the fundamental niche. Ideas in Ecology and Evolution, 5.

79. Rowles, A.D. & O’Dowd, D.J. (2009). New mutualism for old: Indirect disruption and direct facilitation of seed dispersal following Argentine ant invasion. Oecologia, 158, 709–716.

80. Savage, A.M., Rudgers, J.A. & Whitney, K.D. (2009). Elevated dominance of extrafloral nectary-bearing plants is associated with increased abundances of an invasive ant and reduced native ant richness. Divers Distrib, 15, 751–761.

81. Sexton, J.P., McIntyre, P.J., Angert, A.L. & Rice, K.J. (2009). Evolution and ecology of species range limits. Annu Rev Ecol Evol Syst, 40, 415–436.

82. Shaw, A.K., Naven, N. & Stanton, D.E. (2021). Let’s move out together: a framework for the intersections between movement and mutualism. Ecology, 102, e03419.

83. Simberloff, D. & von Holle, B. (1999). Positive Interactions of Nonindigenous Species: Invasional Meltdown? Biological Invasions 1999 *1*:*1*, 1, 21–32.

84. Simonsen, A.K., DInnage, R., Barrett, L.G., Prober, S.M. & Thrall, P.H. (2017). Symbiosis limits establishment of legumes outside their native range at a global scale. Nat Commun, 8, 1–9.

85. Soudzilovskaia, N.A., Vaessen, S., Barcelo, M., He, J., Rahimlou, S., Abarenkov, K., et al. (2020). FungalRoot: global online database of plant mycorrhizal associations. New Phytologist, 227, 955–966.

86. Spengler, R.N. (2020). Anthropogenic Seed Dispersal: Rethinking the Origins of Plant Domestication. Trends Plant Sci, 25, 340–348.

87. Stuble, K.L., Kirkman, L.K. & Carroll, C.R. (2010). Are red imported fire ants facilitators of native seed dispersal? Biol Invasions, 12, 1661–1669.

88. Traveset, A. & Richardson, D.M. (2014). Mutualistic interactions and biological invasions. Annu Rev Ecol Evol Syst, 45, 89–113.

89. Volp, T.M., Cernusak, L.A. & Lach, L. (2022). Epiphytic ant-plant obtains nitrogen from both native and invasive ant inhabitants. Biotropica, 54, 556–560.

90. Warren, R.J., Giladi, I. & Bradford, M.A. (2010). Ant-mediated seed dispersal does not facilitate niche expansion. Journal of Ecology, 98, 1178–1185.

91. Weber, M.G. & Agrawal, A.A. (2014). Defense mutualisms enhance plant diversification. Proc Natl Acad Sci U S A, 111, 16442–16447.

92. Weber, M.G., Porturas, L.D. & K.H., K. (2015). World list of plants with extrafloral nectaries. Available at: http://www.extrafloralnectaries.org/. Last accessed 8 November 2021.

93. Werner, G.D.A., Cornwell, W.K., Sprent, J.I., Kattge, J. & Kiers, E.T. (2014). A single evolutionary innovation drives the deep evolution of symbiotic N2-fixation in angiosperms. Nat Commun, 5, 1–9.

94. Wong, M.K.L., Economo, E.P. & Guénard, B. (2023). The global spread and invasion capacities of alien ants. Current Biology, 33, 566–571.e3.

95. Zeng, Y. & Wiens, J.J. (2021). Species interactions have predictable impacts on diversification. Ecol Lett, 24, 239–248.

