## Supplements for "Generalized mutualisms promote range expansion in both plant and ant partners"

### Supporting Information

### Supplementary Methods

*PGLS models.* For the legume PGLS models, we pruned our range and trait dataset to include only species that were in the Zanne et al. (2014) time-calibrated molecular phylogeny of angiosperm species, resulting in a dataset with 1220 legume species with EFN, domatia, and nodulation trait data, and 384 legume species with data on AM and EM fungal associations. For the ant PGLS models, we pruned our range and trait dataset to include only species in the Nelsen et al. (2018) ant molecular phylogeny, obtaining a final dataset consisting of 769 ant species. We used the *gls* function (Pinheiro *et al.* 2021) to fit PGLS models in R with Pagel's lambda type correlations (*corPagel*) fitted using maximum likelihood. We fit two PGLS models, one for the log-transformed number of non-contiguous ranges for legumes (including zeros for non-introduced legumes) and one for the same variable for ants (again, including zeros for non-introduced ants). For the PGLS models for native range size, we used square root transformed native range areas.

### Supplementary Results

*PGLS models.* For legumes, PGLS models of the number of introduced ranges estimated moderate $\lambda$ values (0.295 for the model with EFNs, domatia, and nodules, and 0.351 for the model with AM and EM fungi), implying modest, but not zero phylogenetic signal (Tables S6 & S8). In these models, EFNs again significantly and strongly increased a legume's number of introduced ranges, while all other mutualistic traits had non-significant effects (Tables S6 & S8), although the PGLS models have reduced power to detect effects because of smaller sample sizes. Phylogenetic signal was stronger for native range area ($\lambda$ = 0.355, Table S7) for the dataset with EFNs, domatia and nodule data, but weaker for the smaller dataset with mycorrhizae data ($\lambda$ = 0.184, Table S9) than for the number of introduced ranges. In contrast to the mixed model of native range area, there was no significant interaction between EFNs and domatia (Table S4). However, only one legume in the PGLS dataset, *Acacia collinsii,* had both EFNs and domatia.

Phylogenetic signal for both introduced ($\lambda = 0.666$) and native ($\lambda = 0.328$) range area was higher in ants than in legumes (compare Tables S6-S8 to Tables S10&S11), indicating greater phylogenetic conservation of range size in ants than in legumes. In the ant PGLS models, visiting EFNs and dispersing seeds, but not nesting in domatia, significantly increased total introduced area and total native area. Tropical ants (i.e., ants with a lower midpoint latitude for their native range) had larger introduced range areas than temperate ants, but latitude did not predict native range area in the PGLS models. Again, total native range area was significantly associated with total introduced range area in the PGLS model, as in the mixed models.

*Multiple mutualisms.* All legumes with four mutualisms had nodules, AM fungi, EM fungi, and EFNs (and no domatia). Most legumes with three mutualisms had either nodules and AM and EM fungi (52 taxa) or nodules and AM fungi and EFNs (63 taxa, out of 121 taxa with three mutualisms). Most legumes with two mutualisms (430/473 taxa) had nodules and AM fungi. Finally, most legumes with only one mutualism type had nodules only (28 taxa) or AM fungi only (63 taxa, out of 111 taxa with one mutualism).

*Native range size.* Legumes with more human uses had larger native range areas (Table S2). In the larger dataset only, temperate species were predicted to have significantly smaller native range areas than tropical species (Table S2). There was no significant difference between annuals and perennials in native range area in the larger dataset, but perennials had larger native range areas than annuals in the smaller dataset on mycorrhizal associations (Table S2). In both datasets, woody legumes had significantly smaller native range sizes than herbaceous legumes. In ants, temperate species had significantly larger native range areas than tropical species (Table S4), as expected from Rapoport's Rule.

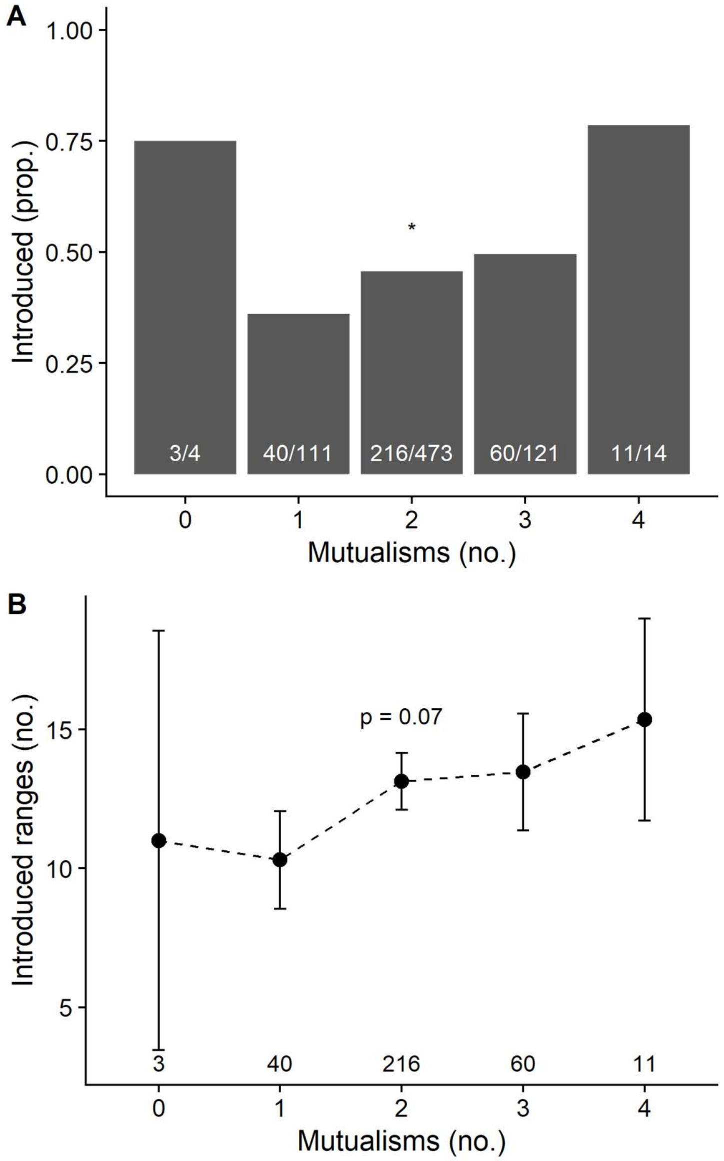


**Figure S1.** Relationship between a legume's number of mutualisms and the proportion of introduced species (top) or number of introduced ranges (bottom, mean ± 1SE). Above the x-axis is the proportion of introduced taxa/total taxa (top) or number of introduced species (bottom) in each category. Significant effects indicated by *** for $p \leq 0.001$, ** for $p \leq 0.01$, and * for $p \leq 0.05$. See Table S1 for complete results of statistical models.

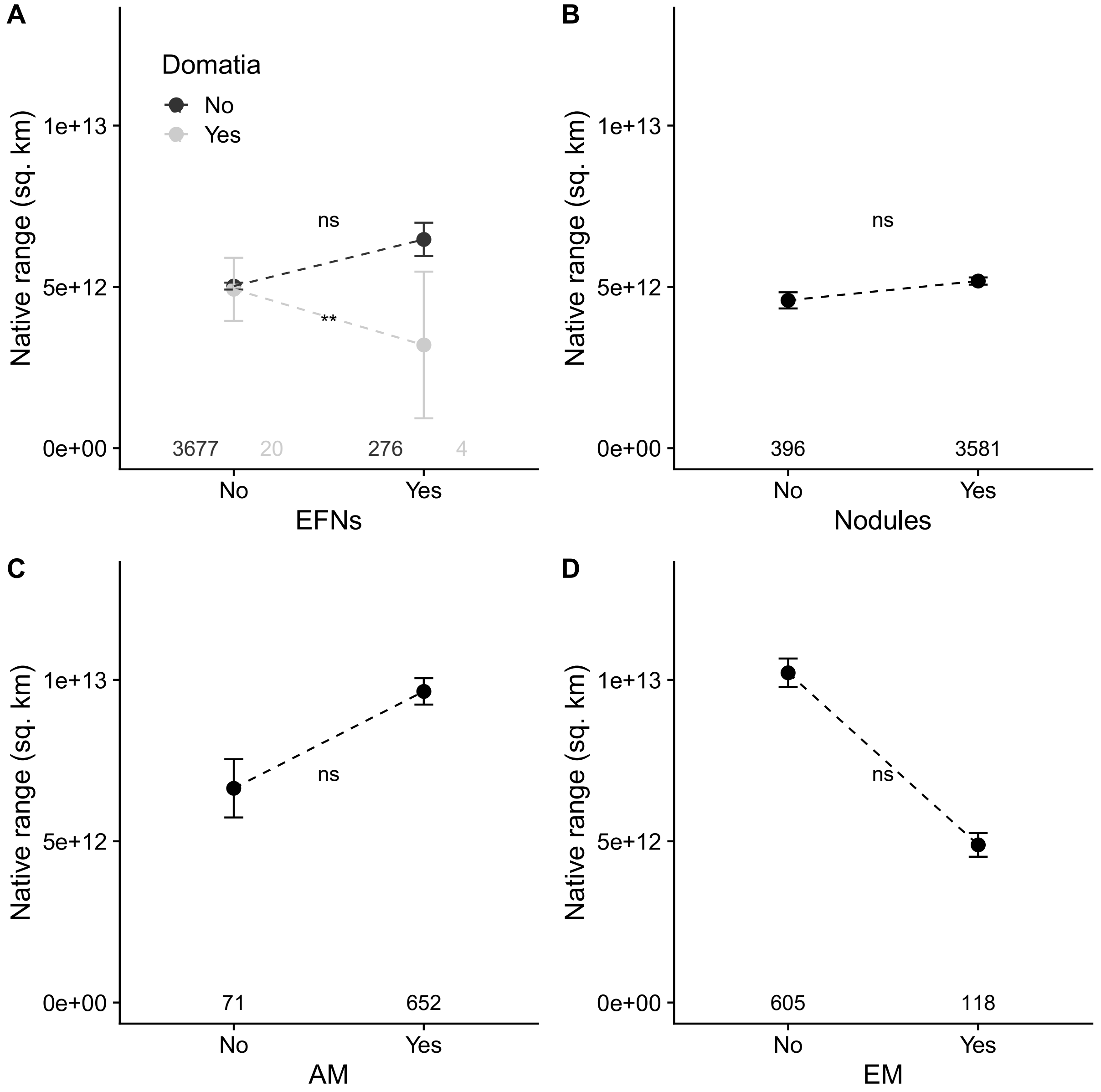


**Figure S2.** Relationship between the presence or absence of EFNs and domatia (A), nodules (B), arbuscular mycorrhizae (C), and ectomycorrhizae (D) and a legume's native range size (mean ± 1SE). The number of legume species in each category is above the x-axis. Significant effects indicated by *** for $p \leq 0.001$, ** for $p \leq 0.01$, and * for $p \leq 0.05$. See Table S2 for complete results of statistical models.

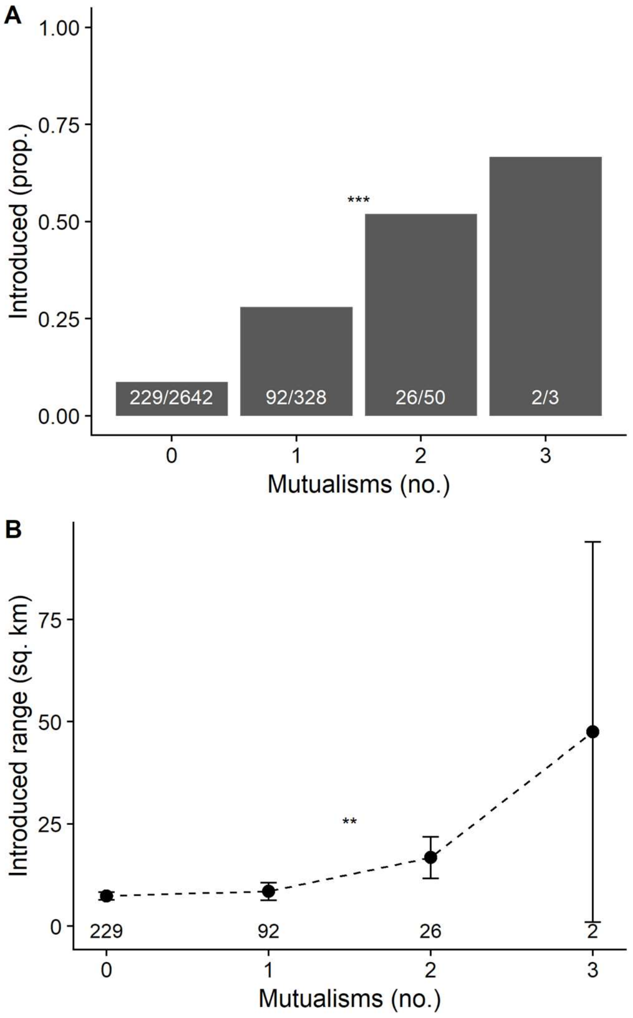


**Figure S3.** Relationship between an ant's number of mutualisms and the proportion of introduced species (top) or number of introduced ranges (bottom, mean ± 1SE). Above the x-axis is the proportion of introduced taxa/total taxa (top) or number of introduced species (bottom) in each category. Significant effects indicated by *** for $p \leq 0.001$, ** for $p \leq 0.01$, and * for $p \leq 0.05$. See Table S5 for complete results of statistical models.

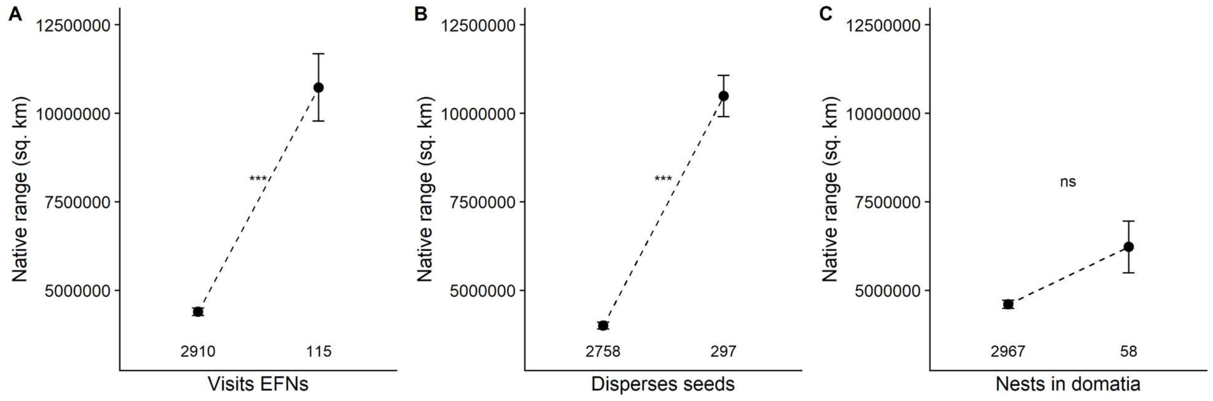


**Figure S4.** Relationship between ants visiting EFNs, dispersing seeds, and nesting in domatia and an ant's native range area (mean ± 1SE). The number of ant species in each category is above the x-axis. Significant effects indicated by *** for $p \leq 0.001$, ** for $p \leq 0.01$, and * for $p \leq 0.05$. See Table S4 for complete results of statistical models.

**Supplementary Tables**

**Table S1.** Model results for the effect of the number of mutualisms (range: 0-4) on whether legumes are successfully introduced to any novel range (left columns), and if successfully introduced, how many introduced ranges they spread to (right columns). Significant effects *italicized* and indicated by *** for $p \leq 0.001$, ** for $p \leq 0.01$, and * for $p \leq 0.05$.

| | **Successful introduction? (Binomial GLMM)** | | **Number of introduced ranges (Gaussian LMM)** | |
|---|---|---|---|---|
| **Fixed effects** | **Estimate** | **SE** | **Estimate** | **SE** |
| Intercept | *-3.583**** | *0.564* | *0.667*** | *0.258* |
| Number of mutualisms | *0.494** | *0.207* | 0.182 | 0.100 |
| Total native area | 0.223 | 0.146 | *-0.186*** | *0.063* |
| Annual? | 0.727 | 0.515 | 0.174 | 0.233 |
| Woody? | *0.862** | *0.359* | *-0.384** | *0.165* |
| Latitude | 0.188 | 0.151 | -0.135 | 0.070 |
| Number of human uses | *0.956**** | *0.088* | *0.316**** | *0.025* |
| Publication count | 0.180 | 0.129 | -0.032 | 0.055 |
| **Random effect** | **Variance** | | **Variance** | |
| Tribe | 0.802 | | 0.114 | |

**Table S2.** Model results for the effects of A. EFNs, domatia, and nodules and B. AM and EM fungi on legume native range size. Significant effects *italicized* and indicated by *** for $p \leq 0.001$, ** for $p \leq 0.01$, and * for $p \leq 0.05$.

| **A.** | **Native range size (Gaussian LMM)** | |
|---|---|---|
| **Fixed effects** | **Estimate** | **SE** |
| Intercept | *14.91**** | *0.113* |
| EFNs? | 0.002 | 0.090 |
| Domatia? | 0.544 | 0.301 |
| Nodules? | -0.093 | 0.097 |
| Annual? | -0.079 | 0.074 |
| Woody? | *-0.329**** | *0.059* |
| Latitude | *-0.181**** | *0.028* |
| Number of human uses | *0.281**** | *0.014* |
| EFNs*Domatia | *-2.220*** | *0.678* |
| Publication count | 0.007 | 0.020 |
| **Random effect** | **Variance** | |
| Tribe | 0.114 | |

| **B.** | **Native range size (Gaussian LMM)** | |
|---|---|---|
| **Fixed effects** | **Estimate** | **SE** |
| Intercept | *15.599**** | *0.190* |
| AM? | 0.065 | 0.150 |
| EM? | 0.150 | 0.141 |
| Annual? | *-0.681**** | *0.185* |
| Woody? | *-0.617**** | *0.131* |
| Latitude | -0.029 | 0.055 |
| Number of human uses | *0.166**** | *0.019* |

| | | |
|---|---|---|
| Study count | -0.015 | 0.041 |
| **Random effect** | **Variance** | |
| Tribe | 0.094 | |

**Table S3.** Model results for the effect of the number of mutualisms (range: 0-3) on whether ants are successfully introduced to any novel range (left columns), and if successfully introduced, how many introduced ranges they spread to (right columns). Significant effects *italicized* and indicated by *** for $p \leq 0.001$, ** for $p \leq 0.01$, and * for $p \leq 0.05$.

| | **Successful introduction? (Binomial GLMM)** | | **Number of introduced ranges (Gaussian LMM)** | |
|---|---|---|---|---|
| **Fixed effects** | **Estimate** | **SE** | **Estimate** | **SE** |
| Intercept | *-2.970**** | *0.186* | *0.806**** | *0.156* |
| Number of mutualisms | *0.973**** | *0.133* | *0.307*** | *0.105* |
| Total native area | *0.412**** | *0.037* | *0.071**** | *0.062* |
| Latitude | *-0.222**** | *0.062* | -0.398 | 0.063 |
| **Random effect** | **Variance** | **SE** | **Variance** | **SE** |
| Tribe | 0.6592 | 0.015 | 0.431 | 0.036 |

**Table S4.** Model results for the effects of visiting EFNs, dispersing seeds, and nesting in domatia on ant native range area. Interactions among mutualistic traits were never significant and omitted from final models. Significant effects *italicized* and indicated by *** for $p \leq 0.001$, ** for $p \leq 0.01$, and * for $p \leq 0.05$.

| | **Native range size (Gaussian LMM)** | |
|---|---|---|
| **Fixed effects** | **Estimate** | **SE** |
| Intercept | *14.132**** | *0.112* |
| Visits EFNs? | *0.640**** | *0.169* |
| Nests in domatia? | 0.438 | 0.264 |
| Disperses seeds? | *1.297**** | *0.113* |
| Latitude | *0.103*** | *0.031* |
| **Random effect** | **Variance** | **SE** |
| Tribe | 0.465 | |

**Table S5.** Model results for the effects of visiting EFNs, dispersing seeds, and nesting in domatia on the likelihood an ant species is intercepted at customs (left two columns), establishes indoors (middle two columns), or becomes naturalized outdoors (right two columns). Interactions among mutualistic traits were never significant and omitted from final models. Significant effects *italicized* and indicated by *** for $p \leq 0.001$, ** for $p \leq 0.01$, * for $p \leq 0.05$, and † for $p \leq 0.1$.

| | **Intercepted? (Binomial GLMM)** | | **Established indoors? (Binomial GLMM)** | | **Naturalized outdoors? (Binomial GLMM)** | |
|---|---|---|---|---|---|---|
| **Fixed effects** | **Estimate** | **SE** | **Estimate** | **SE** | **Estimate** | **SE** |
| Intercept | -0.076 | 0.199 | *-1.076**** | *0.255* | *0.548** | *0.254* |
| Visits EFNs? | 0.473 | 0.438 | *1.037*** | *0.400* | 0.608 | 0.412 |
| Nests in domatia? | 15.834 | 295.603 | -0.255 | 1.000 | -0.501 | 0.937 |
| Disperses seeds? | *1.154**** | *0.336* | *-1.042*** | *0.381* | -0.138 | 0.323 |
| Total native area | 0.154 | 0.133 | *0.279** | *0.120* | -0.018 | 0.124 |
| Latitude | *-0.407*** | *0.132* | -0.224 | 0.139 | *-0.277** | *0.132* |
| **Random effect** | **Variance** | | **Variance** | | **Variance** | |
| Tribe | 0.334 | | 0.503 | | 0.808 | |

**Table S6.** PGLS results for the effects of EFNs, domatia, and nodules on the number of introduced ranges for each legume species. Significant effects *italicized* and indicated by *** for $p \leq 0.001$, ** for $p \leq 0.01$ and * for $p \leq 0.05$.

| **Lambda** | 0.298 | |
|---|---|---|
| **Predictors** | **Estimate** | **SE** |
| **Intercept** | 0.145 | 0.198 |
| **EFNs?** | *0.466**** | *0.074* |
| **Nodules?** | 0.108 | 0.115 |
| **Domatia?** | -0.274 | 0.311 |
| **Total native area** | 0 | 0 |
| **Latitude** | -0.002 | 0.002 |
| **Number of human uses** | 0.416*** | 0.013 |
| **Woodiness** | 0.091 | 0.069 |
| **Annual?** | 0.09 | 0.072 |

**Table S7.** PGLS results for the effects of EFNs, domatia, and nodules on legume native range. Significant effects *italicized* and indicated by *** for $p \leq 0.001$, ** for $p \leq 0.01$ and * for $p \leq 0.05$.

| **Lambda** | 0.351 | |
|---|---|---|
| **Predictors** | **Estimate** | **SE** |
| **Intercept** | *2.578** | *1.076* |
| **EFNs?** | 0.262 | 0.380 |
| **Domatia?** | 0.358 | 1.612 |
| **Nodules?** | 0.777 | 0.609 |
| **Latitude** | *-0.464*** | *0.153* |
| **Annual?** | *-1.480**** | *0.371* |
| **Woodiness** | *-1.319**** | *0.360* |
| **Publication count** | 0.060 | 0.107 |
| **Number of human uses** | *0.960**** | *0.058* |

**Table S8.** PGLS results for the effects of AM and EM fungi on the number of introduced ranges for each legume species. Significant effects *italicized* and indicated by *** for $p \leq 0.001$, ** for $p \leq 0.01$ and * for $p \leq 0.05$.

| **Lambda** | 0.355 | |
|---|---|---|
| **Predictors** | **Estimate** | **SE** |
| **Intercept** | 0.333 | 0.405 |
| **AM** | 0.027 | 0.173 |
| **EM** | 0.133 | 0.151 |
| **Native area** | *-0.139*** | *0.000* |
| **Latitude** | *-0.154** | *0.004* |
| **Annual?** | 0.242 | 0.187 |
| **Woody?** | -0.132 | 0.157 |
| **Publication count** | -0.021 | 0.029 |
| **Number of human uses** | *0.423**** | *0.021* |

**Table S9.** PGLS results for the effects of AM and EM fungi on native legume range. Significant effects *italicized* and indicated by *** for $p \leq 0.001$, ** for $p \leq 0.01$ and * for $p \leq 0.05$.

| **Lambda** | 0.184 | |
|---|---|---|
| **Predictors** | **Estimate** | **SE** |
| **Intercept** | *3.038**** | *0.429* |
| **AM** | 0.100 | 0.273 |
| **EM** | -0.135 | 0.237 |
| **Latitude** | -0.008 | 0.099 |
| **Annual?** | *-1.235**** | *0.290* |
| **Woodiness** | *-0.836**** | *0.235* |
| **Publication count** | -0.060 | 0.072 |
| **Number of human uses** | *0.203**** | *0.031* |

**Table S10.** PGLS results for the effects of visiting EFNs, residing in domatia and dispersing seeds on the number of introduced ranges of ants. Significant effects *italicized* and indicated by *** for $p \leq 0.001$, ** for $p \leq 0.01$ and * for $p \leq 0.05$.

| **Lambda** | 0.666 | |
|---|---|---|
| **Predictors** | **Estimate** | **SE** |
| **Intercept** | 0.275 | 0.293 |
| **Visit EFNs?** | *0.474**** | *0.103* |
| **Reside in domatia?** | 0.022 | 0.152 |
| **Disperse seeds?** | *0.295**** | *0.079* |
| **Native area** | *0.000**** | *0.000* |
| **Latitude** | *-0.014**** | *0.003* |

**Table S11.** PGLS results for the effects of visiting EFNs, residing in domatia and dispersing seeds on the native range of ants. Significant effects *italicized* and indicated by *** for $p \leq 0.001$, ** for $p \leq 0.01$ and * for $p \leq 0.05$.

| **Lambda** | 0.328 | |
|---|---|---|
| **Predictors** | **Estimate** | **SE** |
| **Intercept** | *1.829**** | *0.271* |
| **Visit EFNs?** | *0.753**** | *0.154* |
| **Reside in domatia?** | -0.151 | 0.225 |
| **Disperse seeds?** | *1.008**** | *0.116* |
| **Latitude** | 0.005 | 0.004 |
